# An Interactive Brain Atlas of Knowledge

**DOI:** 10.64898/2025.12.12.694043

**Authors:** Leon Stefanovski, Konstantin Bülau, Leon Martin, Christoph Hüttl, Chloê Langford, Jessica Palmer, Marc Sacks, Lion Deger, Marius Pille, Michael Schirner, Jil Meier, Clemens Neudorfer, Andreas Horn, Ana Solodkin, Bertrand Thirion, Martin Hofmann-Apitius, Marc Jacobs, Alpha Tom Kodamullil, the Alzheimer’s Disease Neuroimaging Initiative, Petra Ritter

## Abstract

Biomedical knowledge about the brain increases every day, with a rapidly growing number of scientific publications, datasets, and software tools. While this informational plethora is not merely comprehensible by human beings, recent developments in information science and computational linguistics aim to make this knowledge programmatically accessible by literature mining. However, integrating these semantic methods into neuroimaging standards remains insufficient, hindering researchers from unraveling their full potential.

Therefore, we developed the semantic meta-analysis platform The Virtual Brain adapter of semantics (*TVBase)* that projects biomedical knowledge preserved in over 36 million scientific articles onto a 3D standardized brain. The literature-mining platform *SCAIView* was used to extract ontologically defined biomedical entities and their associations with brain anatomy from the *PubMed* database. By querying a specific concept, the association strength with each anatomical term was calculated using entropy. To project the data onto a standardized brain, we created a unique transformation matrix that links over 800 anatomical terms to voxel coordinates of a parcellated standard brain.

This novel method of knowledge projection extracts region-specific information about biomedical concepts from the literature to support translational multi-scale approaches to computational neuroscience. The multi-purpose software framework *TVBase* is openly available as a Python library. It aims for hypothesis-free neuroimaging pattern interpretation, hypothesis generation, and applications in personalized medicine.

## 1 INTRODUCTION

The multitude of data and knowledge in science is growing daily in quantities beyond human grasp. This trend leads to the necessity of powerful computational tools for their comprehensive analysis and integrative understanding. In 2024, 1.7 million biomedical articles were published solely on *PubMed* (National Library of Medicine 2022)– with more than 20,000 articles just on “Alzheimer’s”. Consequently, data and knowledge are increasingly available in the structured and machine-readable form of so-called knowledge graphs (Ji, Pan et al. 2021). There is no doubt in the scientific community that a holistic integration of all these knowledge sources would be beneficial to overcome current research needs (Stephan, Binder et al. 2016, Sy, Roman et al. 2023) – but the question is, how can this be done without losing the connection to practical research problems and real-world experiments?

The field of neuroscience is genuinely interested in using data- and knowledge-driven approaches to overcome the replication crisis (Ioannidis 2005). In addition, particular characteristics of neuroscience experiments typically lead to small sample sizes, expensive designs, and complex data output (Alger 2022, Calin-Jageman 2022). While a careful evaluation of the way we store and share data is an essential part of the solution (Gorgolewski, Auer et al. 2016, Klingner, Denker et al. 2022, Schirner, Domide et al. 2022), the direct metanalysis of existing data and knowledge holds the potential to derive new results from established material. The case of Alzheimer’s Disease (AD) represents a prototypical example. A plethora of knowledge accumulated during more than a hundred years of research after the initial description of the disease (Alzheimer 1907). Nevertheless, there is still a lack of understanding of the disease mechanisms. The question about the “cause of AD” is still controversial and interdisciplinary and would benefit from approaches that integrate knowledge from various scales and disciplines, e.g., genetics, neuroimaging, biochemistry, psychology, and clinical medicine (Stefanovski, Meier et al. 2021).

Recent initiatives aim to build atlases or catalogues of available brain data by mapping various modalities of measurements onto a brain template and hence anchoring the available information to standardized brain coordinates. The tool *neuromaps* has integrated common brain imaging modalities into 66 brain maps (Markello, Hansen et al. 2022), aiming to allocate human neuroimaging data in a comprehensive context of the known underlying structure and function (Voytek 2022). Automated neuroimaging metanalyses go one step further by linking published coordinate-based maps to the text describing it in the publication (Yarkoni, Poldrack et al. 2011, Dockès, Poldrack et al. 2020, Beam, Potts et al. 2021). As they provide maps of activation patterns for various queries, these tools combine the strength of a metanalysis with the utility of published datasets and the ease of a search engine. However, while these tools provide data-driven solutions, they neglect large parts of the knowledge available in semantic form (i.e., literature), which is not derived from neuroimaging data.

The recent advances in natural language processing enable next-generation chatbot systems and automatic text-writing (Saravanan and Sudha 2022) and allow for high-end text mining to make the knowledge from the scientific literature available in a computational form. The idea behind this goes back to the origins of the internet in the early 2000s, but it is still a modern desideratum: the semantic web (Berners-Lee, Hendler et al. 2001). The underlying idea is disruptive: to enhance the internet by a semantic layer enriching text with meaning. Concretely, the meaning of a word needs to be annotated with a controlled vocabulary – to highlight, e.g., that “APP” in a particular context is the “amyloid precursor protein” and not the latest mobile “app”. Some of these controlled vocabularies are already widely used, such as e.g., the Medical Subject headings (*MeSH*, (Rogers 1963)). Such structured knowledge systems are investigated in a dedicated field of computational research, the name of which is referring to the ancient science of being – “ontology” - and to formal logic as described in the Socratic era of science, where Plato derives his ontologic theory out of a semantic problem (Plato 360 B.C.). An ontology in informatics is a graph-like representation of knowledge that contains its own inference rules (Sy, Roman et al. 2023). Besides *MeSH*, recent studies showed the potential of the protein-protein interaction database STRING (Szklarczyk, Gable et al. 2021), the systematic gene nomenclature of the HUGO Gene Nomenclature Committee HGNC (Povey, Lovering et al. 2001), and the gene ontology GO (Gene Ontology Consortium 2004), describing biological processes. Former studies (Dörpinghaus, Klein et al. 2018) have used all these controlled vocabularies to extract statistics from the whole literature corpus of *PubMed* (Roberts 2001). This opens the door to knowledge-based complementation and integration of data-driven experiments – if they can be linked for instance by a joint spatial anchoring system, which, in the case of the present study is a standard anatomically defined brain space. We used the anatomical ontology *Uberon* (Mungall, Torniai et al. 2012) to link the semantic knowledge of the whole *PubMed* literature to a standardized brain template. This represents a complementary approach compared to existing metaanalysis tools like *NeuroQuery* (Dockès, Poldrack et al. 2020), where the analysis is grounded in, but also limited to published datasets.

The Virtual Brain adapter of semantics – *TVBase* – is a tool for literature-mining-based semantic brain mapping that creates unique brain maps based on any semantic query, relying on the information from more than 36 million scientific articles in PubMed (National Library of Medicine 2022).

## 2 RESULTS

### 2.1 TVBase – a unique tool for mapping semantic concepts onto the brain

The interactive atlas of knowledge provided by TVBase offers a novel and unique tool that allows to map any semantic concept onto the three-dimensional structure of the brain. While its core functionality can be unveiled by using it as a Python library, the software package offers also a graphical user interface (GUI) and can serve in several translational applications for science and clinics. Its heart element is a detailed transformation between the semantic space of neuroanatomical nomenclature and a standardized three-dimensional brain template. TVBase creates a new form of semantic brain maps, in the following referred to as *TVBase maps*, based on the knowledge contained in the PubMed database. An overview of TVBase is provided in **Figure 1**.

**Figure 1.**
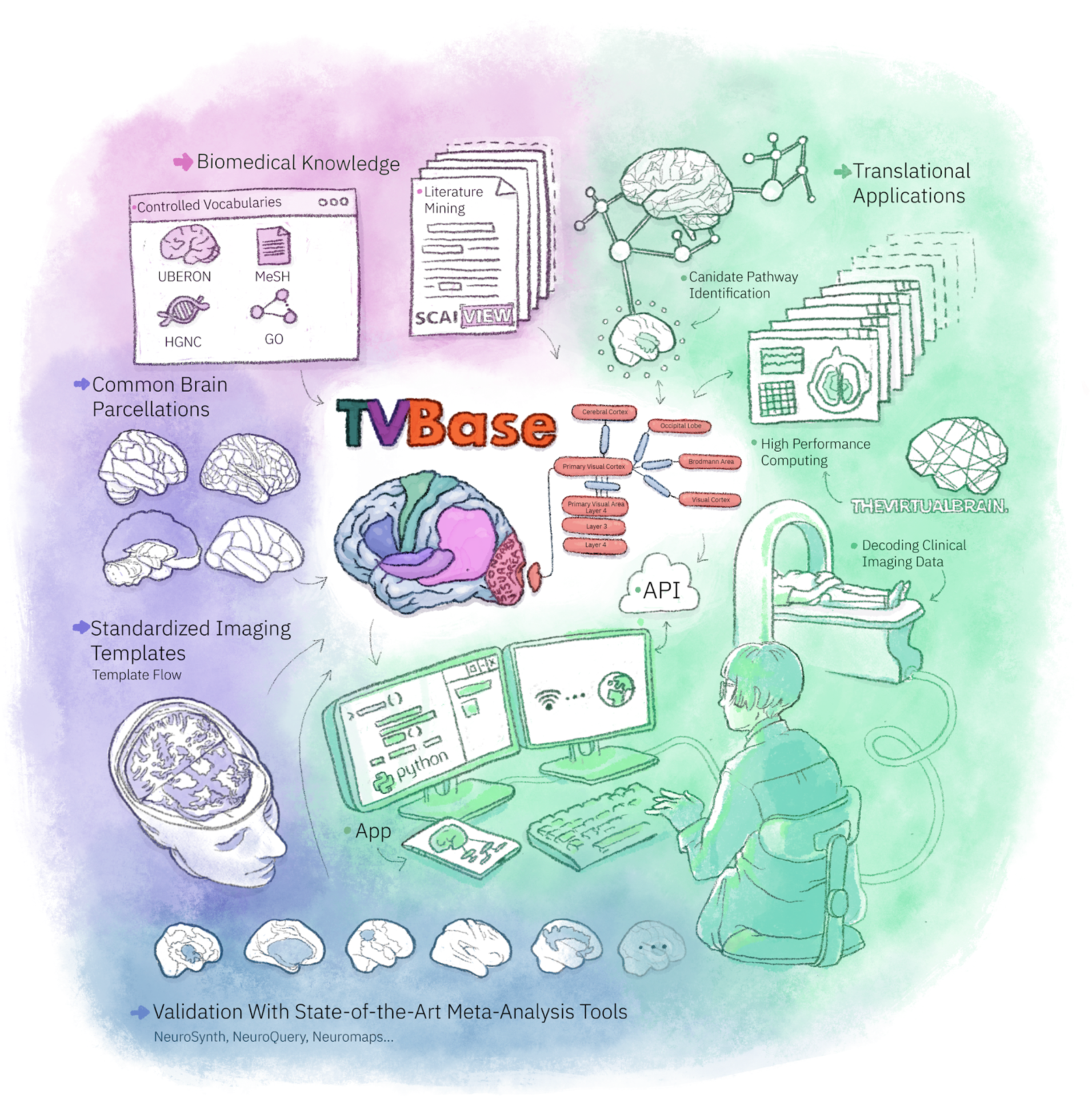
The TVBase framework. We introduce state-of-the-art software that allows mapping any semantic concept for multiple research approaches. It uses the co-occurrence of anatomical terms and the queried concept in the scientific literature, mapping the query onto a brain template. The maps can be accessed using Python functions, API access, or TVBase GUI (both on desktop and mobile engines). The generated TVBase maps show the relevance – a metric for the semantic association of the query and the anatomical region – and are accompanied by extensive metadata describing the underlying evidence. The maps can be used to interpret empirically measured data, to inform complex mechanistic brain models, or to investigate spatial similarities between different maps.

*TVBase* extracts the frequency of a queried concept in the text-mined literature corpus of *SCAIView* (Dörpinghaus, Klein et al. 2018) and the frequency of its co-occurrence with all anatomical terms included in the *Uberon* ontology (Mungall, Torniai et al. 2012) (pink part of **Fig. 1**). We employ information theory here: our tool uses the modified Kullback-Leibler divergence, related to surprise in information theory, as relevance measure for each anatomical term associated with this concept, such as assessing the relevance of the “hippocampal formation” to “memory."

To ensure the highest quality of the anatomical assignment, we performed a multi-step manual curation process to ensure correct linkage of *Uberon* terms with the brain parcellation of the Human connectome project (HCP, (Glasser, Coalson et al. 2016)). The anatomical curation is described in detail in Supplementary A.

*TVBase* can map any brain-related concept, i.e., free-text terms, as well as controlled vocabularies representing genes, proteins, biological processes, and brain functions. The maps are provided in common volumetric template spaces (purple part of **Fig. 1**). It relies on the Montreal Neuroimaging Institute – International Consortium for Brain mapping nonlinear atlases version 2009 (MNI-ICBM152-2009c (Fonov, Evans et al. 2009), further referred to as MNI-152), and average surface triangulations, like the *fsaverage* template (Fischl, Sereno et al. 1999, Fischl 2012), and HCP’s freesurfer-left-right-aligned (fsLR) surface (Glasser, Sotiropoulos et al. 2013). As a consequence of using highly standardized templates, *TVBase* can easily reparcellate each map into more than 46 different brain atlases (Lawrence, Bridgeford et al. 2021).

It is delivered as an open-source software available as a Python package, offers API access, and a database collecting the generated maps and their metadata (green part of **Fig. 1**). It can be further accessed by a Web interface with mobile support and is compatible with the neuroinformatic simulation platform The Virtual Brain (www.thevirtualbrain.org). *TVBase* is publicly available on *GitHub* (https://github.com/BrainModes/tvbase) and *pypi* (https://pypi.org/project/tvbase). The complete documentation can be found at (https://github.io/BrainModes/tvbase).

For rigorous validation, we systematically assessed the quality of 8121 *TVBase*-generated brain maps by comparing them to existing validated knowledge sources (blue part of **Fig. 1**). On average, each of the 8121 mapped concepts is based on 2170 scientific articles describing 345 unique *Uberon* terms of brain anatomy.

In the following sections, we will show that *TVBase* can approximate empirical data from various scales (2.2), contains network characteristics (2.3), can help decode clinical data (2.4) and can help in understanding cause-and-effect pathways (2.5).

### 2.2 *TVBase* generates brain maps for various biomedical concepts – from receptors to symptoms

To evaluate the validity of *TVBase* maps, we compared a selection of maps to their counterpart output by state-of-the-art validated brain mapping data bases (**Figure 2** and **Table 1**). We focused on the modalities that are included in *neuromaps* (Markello, Hansen et al. 2022) and on thousands of predicted activation patterns from *Neurosynth* (Yarkoni, Poldrack et al. 2011) and *NeuroQuery* (Dockès, Poldrack et al. 2020). The similarity between empirical and *TVBase* maps has been assessed using the Pearson correlation between voxel-wise data arrays. (**Table 1**).

**Figure 2.**
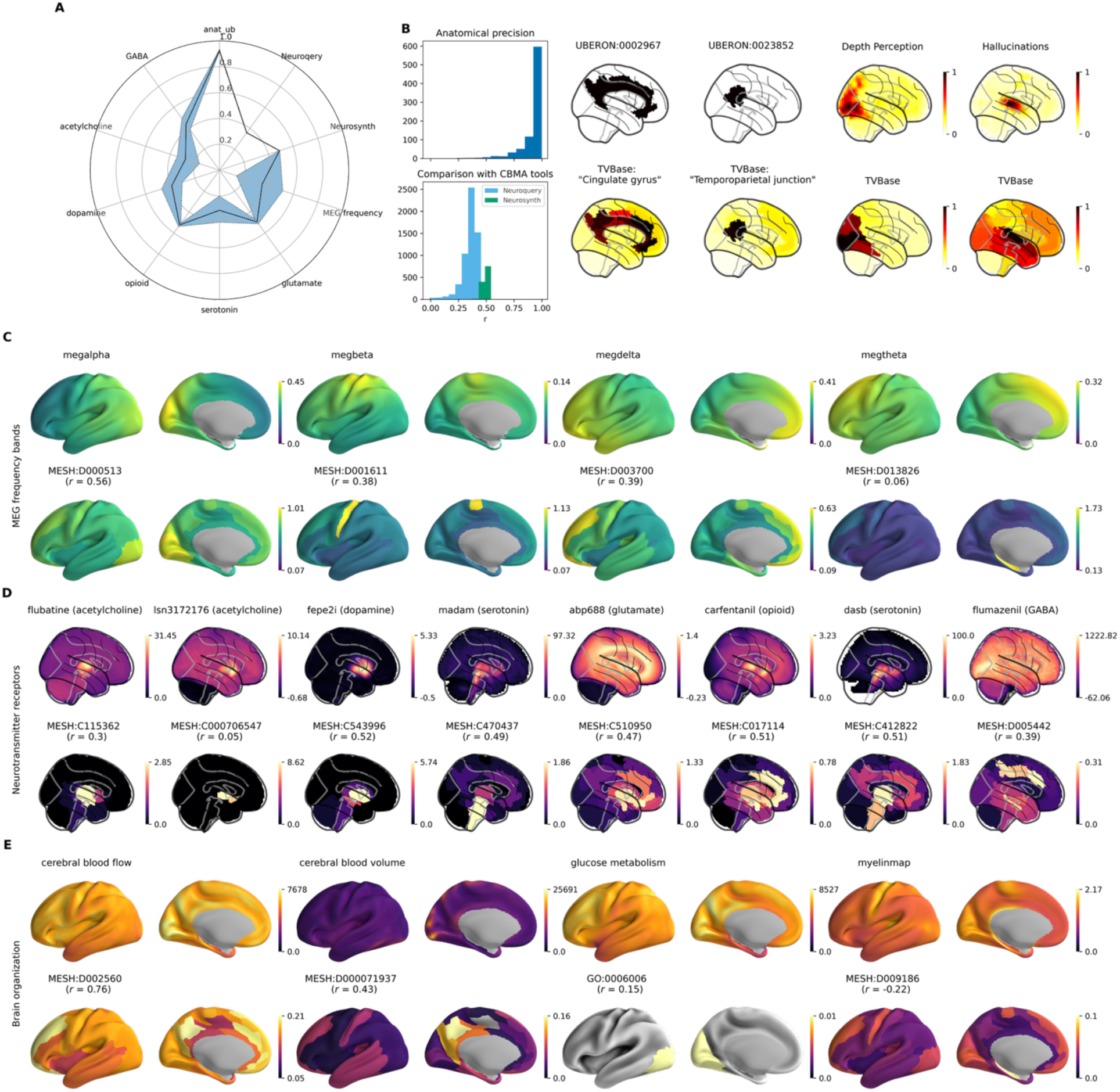
TVBase maps show strong correlation across empirical modalities, ranging from brain function to advanced neuroimaging. Panel A) shows the correlation with different modality domains: anatomical terms, Neuroquery and Neurosynth activation patterns, magnetoencephalography (MEG) frequency bands, and six different neurotransmitter systems. B) We observe the highest correlation for anatomical terms correlated with the anatomical entities in TVBase as a sanity check. Activation-pattern-based maps also significantly overlap across 5921 maps in NeuroQuery and 1291 maps in Neurosynth.). C) Typical frequency band distributions from MEG recordings. Note that we compare TVBase maps here to surface-based data instead of volumetric data. D) Positron emission tomography (PET)-based maps of receptor distributions provide broad evidence of TVBase resembling functional features of the brain by querying the PET tracers in TVBase. E) Further structural (e.g., myelination) and functional (e.g., energy metabolism) features are captured as well by TVBase.

**Table 1.**
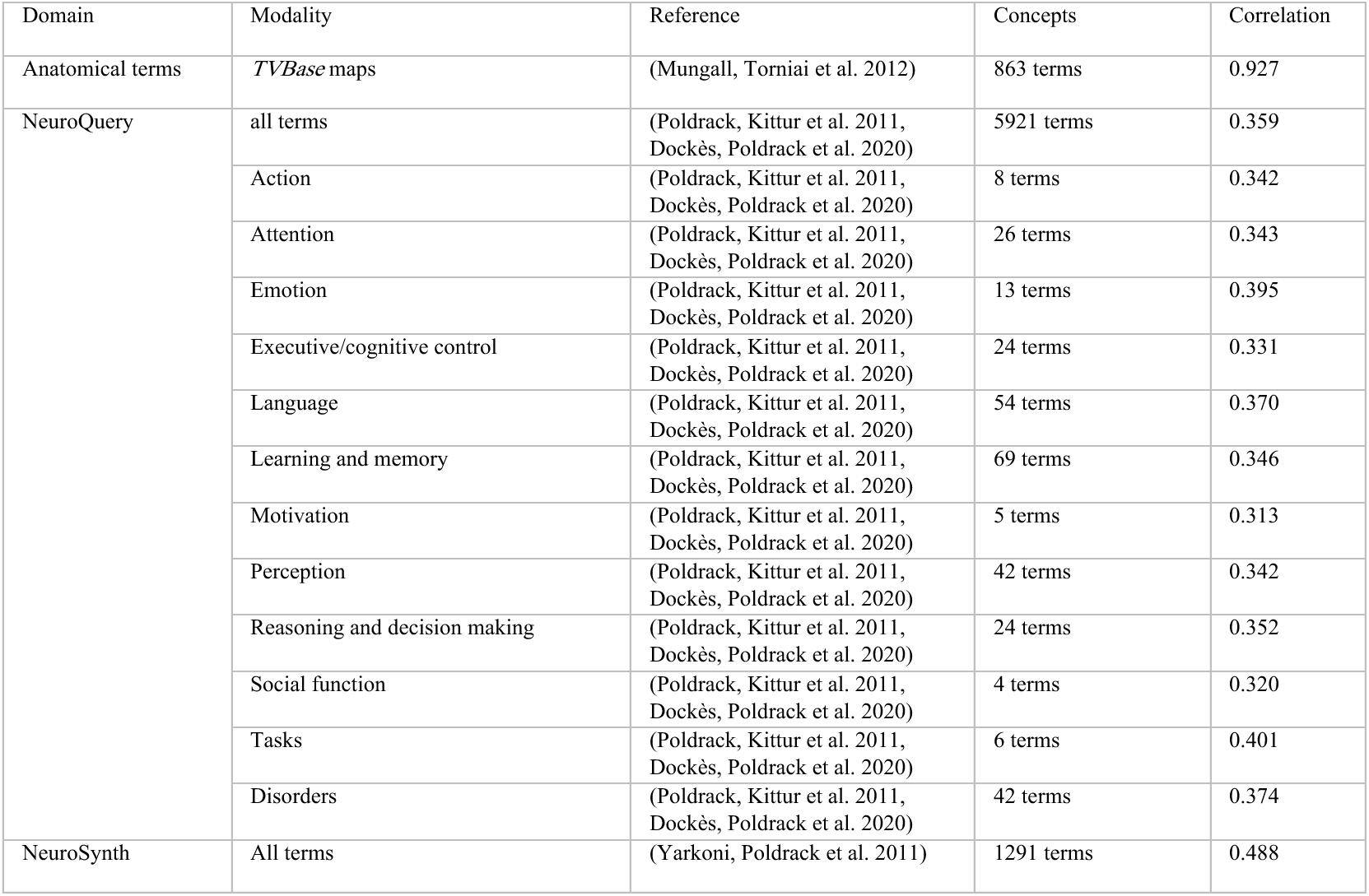
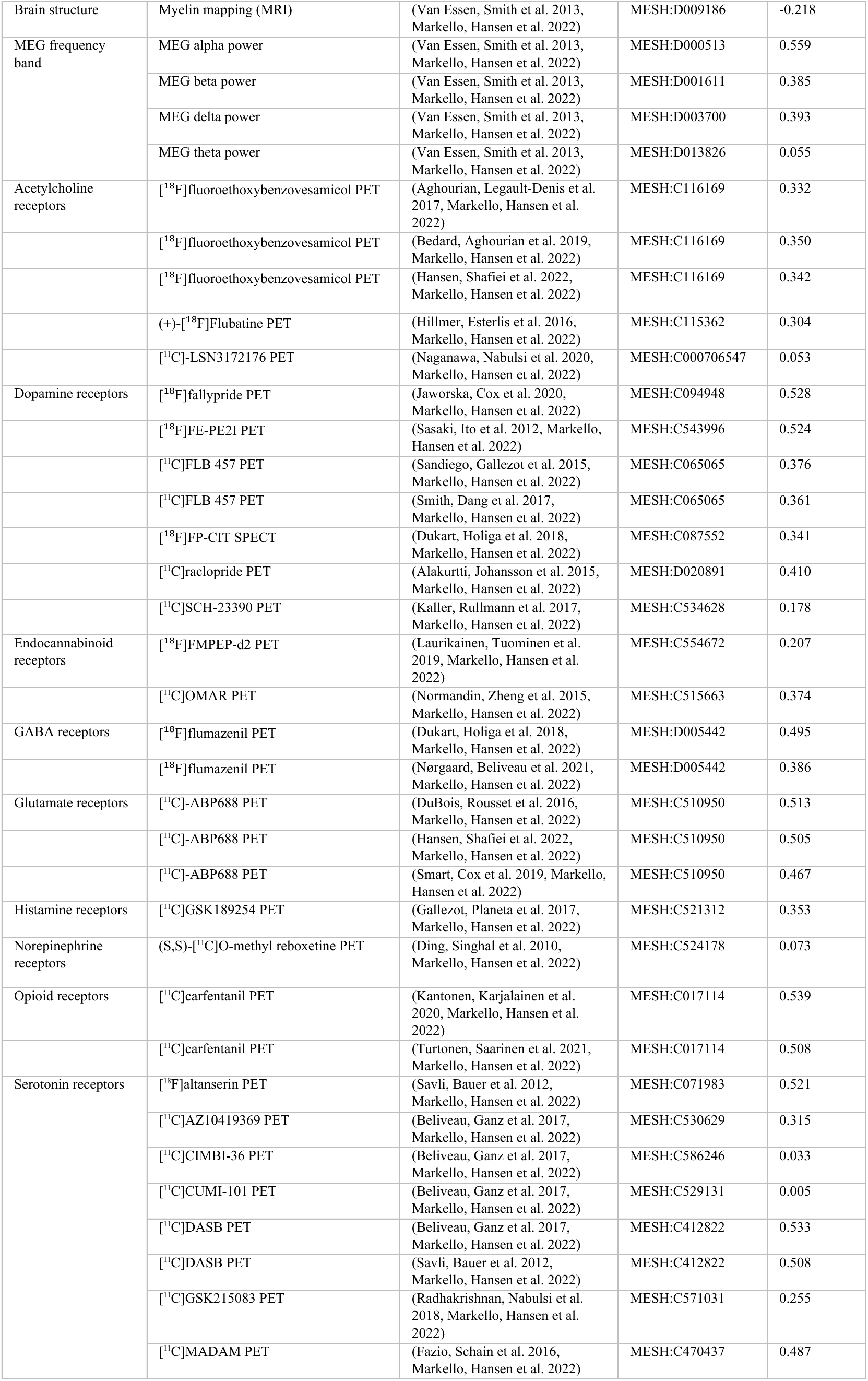
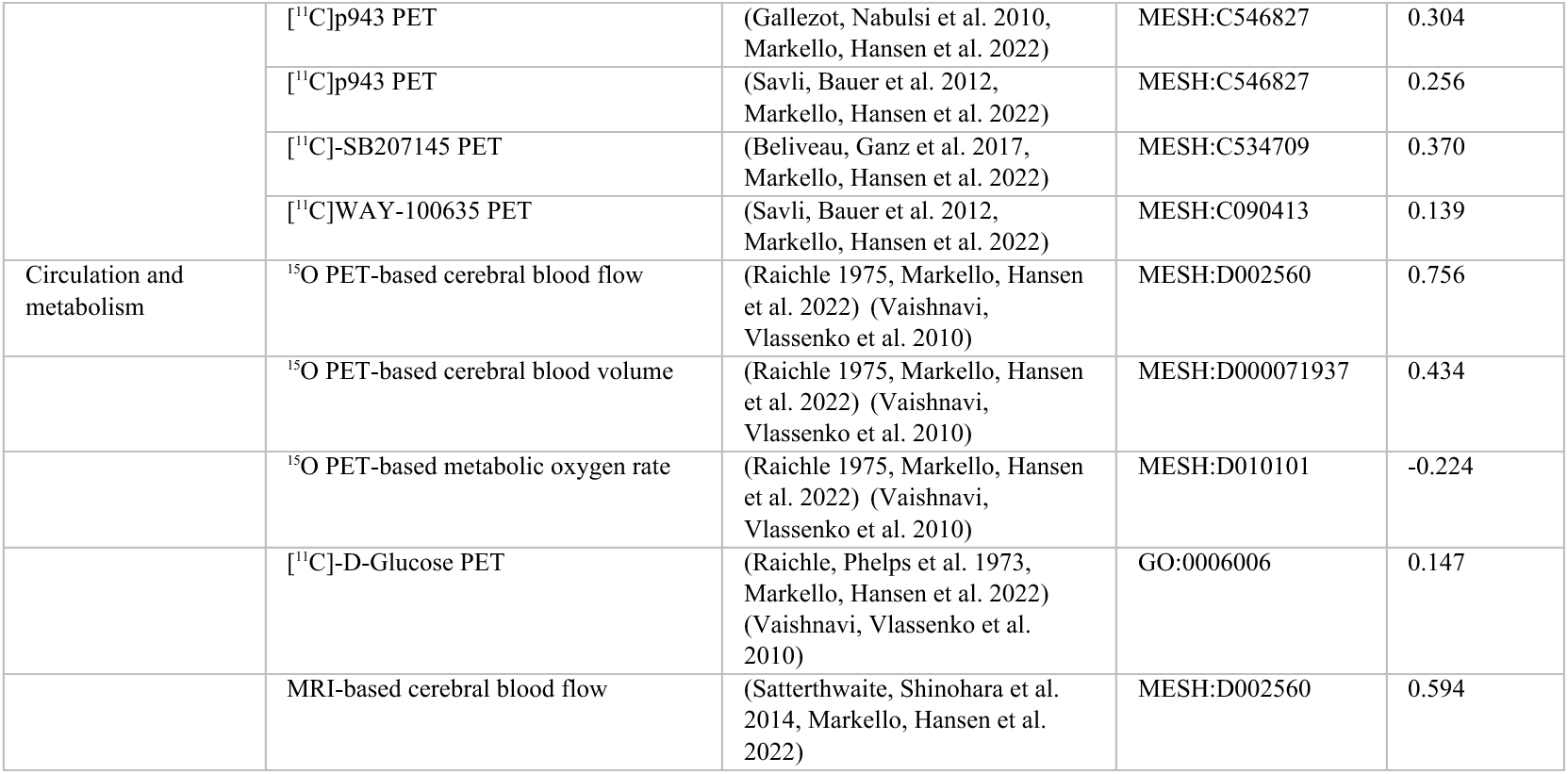
TVBase maps shows medium to strong correlation with empirical maps, ranging from brain function to advanced neuroimaging modalities. We have used standardized maps from neuromaps (Markello, Hansen et al. 2022) and coordinate-based meta-analysis results of functional imaging data retrieved from NeuroQuery (Dockès, Poldrack et al. 2020) and Neurosynth (Yarkoni, Poldrack et al. 2011) to compare them to their corresponding maps in TVBase. The mentioned correlation between the empirical map and the TVBase map is tested for significance using a permutation test with 5000 iterations (the percentile in the last column provides the amount of randomly drawn maps that are more similar than the corresponding TVBase map). Abbreviations: MRI – magnetic resonance imaging, MeSH – Medical Subject Headings, MEG – magnetoencephalography, PET – positron emission tomography, GO – gene ontology; we used the trivial names for chemical compounds.

As expected, a comparison with maps for anatomical queries revealed a strong mean correlation of *r* = 0.927 with our transformation matrix as defined in Supplementary Material A. The coordinate-based tools showed a more complex result: while we still received high and significant correlations of *r =* 0.359 for *NeuroQuery* and 0.488 for *Neurosynth*, we also found out that, for some instances, the maps differ because of their reference to different data foundations. For instance, the *TVBase* map for “hallucinations” shows both acoustical hallucinations in the temporal lobe and visual hallucinations in the occipital lobe, while *NeuroQuery* only captures the (more common) acoustic hallucinations (**Figure 2B**). The concordance between *NeuroQuery* and *TVBase* is higher within domains of the cognitive atlas ontology (Poldrack, Kittur et al. 2011) (**Table 1**).

In the validation with *neuromaps*, we compared results to all modalities that are mappable in a semantic manner (for instance Alpha Rhythm) resulting in 46 maps from 12 different domains (**Table 1**). This revealed a significant positive correlation for 44 maps out of these. The maps include the distribution of electrophysiological frequency bands, positron emission tomography (PET) for nine neurotransmitter systems and metabolic brain activity features. Typical frequency band distributions from MEG (Van Essen, Smith et al. 2013) in *neuromaps* (Markello, Hansen et al. 2022) show strong overlaps for alpha, beta, and delta. For theta, TVBase depicts the hippocampus as an established source of theta activity, which is not accessible via MEG and therefore not observable in empirical data.

PET-based maps of receptor distributions provide broad evidence of TVBase resembling functional features of the brain by querying the PET tracers in TVBase, further structural (e.g., myelination) and functional (e.g., energy metabolism) features can be mapped.

A complete overview of the compared maps, including abbreviations, can be found in Table 1.

### 2.3 Literature-based semantic relevance networks share characteristics with functional and structural connectomes of the brain

As we have demonstrated that TVBase can resemble the spatial patterns of various concepts where we know the ground truth through empirical data, we were keen to explore its capabilities to investigate the interdependence of semantic concepts related to the brain. TVBase offers a novel and unique way to understand how brain regions are organized in terms of their functions and meanings. We studied the relationships between psychological and psychiatric concepts. Using a dataset of 1,186 TVBase maps from the “Psychiatry and Psychology” branch of the MeSH tree, which includes topics like mental disorders, behavior, and psychological phenomena, we looked at how often brain regions appear together in these maps. We then created a matrix showing how strongly different brain regions are connected based on their associations with the same semantic concepts. For example, regions in the occipital cortex are closely linked to concepts like “visual perception,” so they are strongly connected in the matrix. From this, we built a graph we called a semantic relevance network that shows how brain regions are linked through shared meanings. This idea builds on methods used in genetics to study how genes interact (Butte and Kohane 2003) and adapts it to explore connections between brain regions and psychological concepts. It can be seen as a completely new modality, describing functional brain patterns based on semantic associations.

To explore if these semantic relevance networks contain characteristics similar to established patterns in connectomics, we then compared the relevance network with group-based structural connectivity (SC) and functional connectivity (FC) matrices of the S1200 release of the “Young Adult” dataset of the Human Connectome Project (Glasser, Sotiropoulos et al. 2013, Van Essen, Smith et al. 2013). The corresponding matrices are shown in **Figure 3A**.

**Figure 3.**
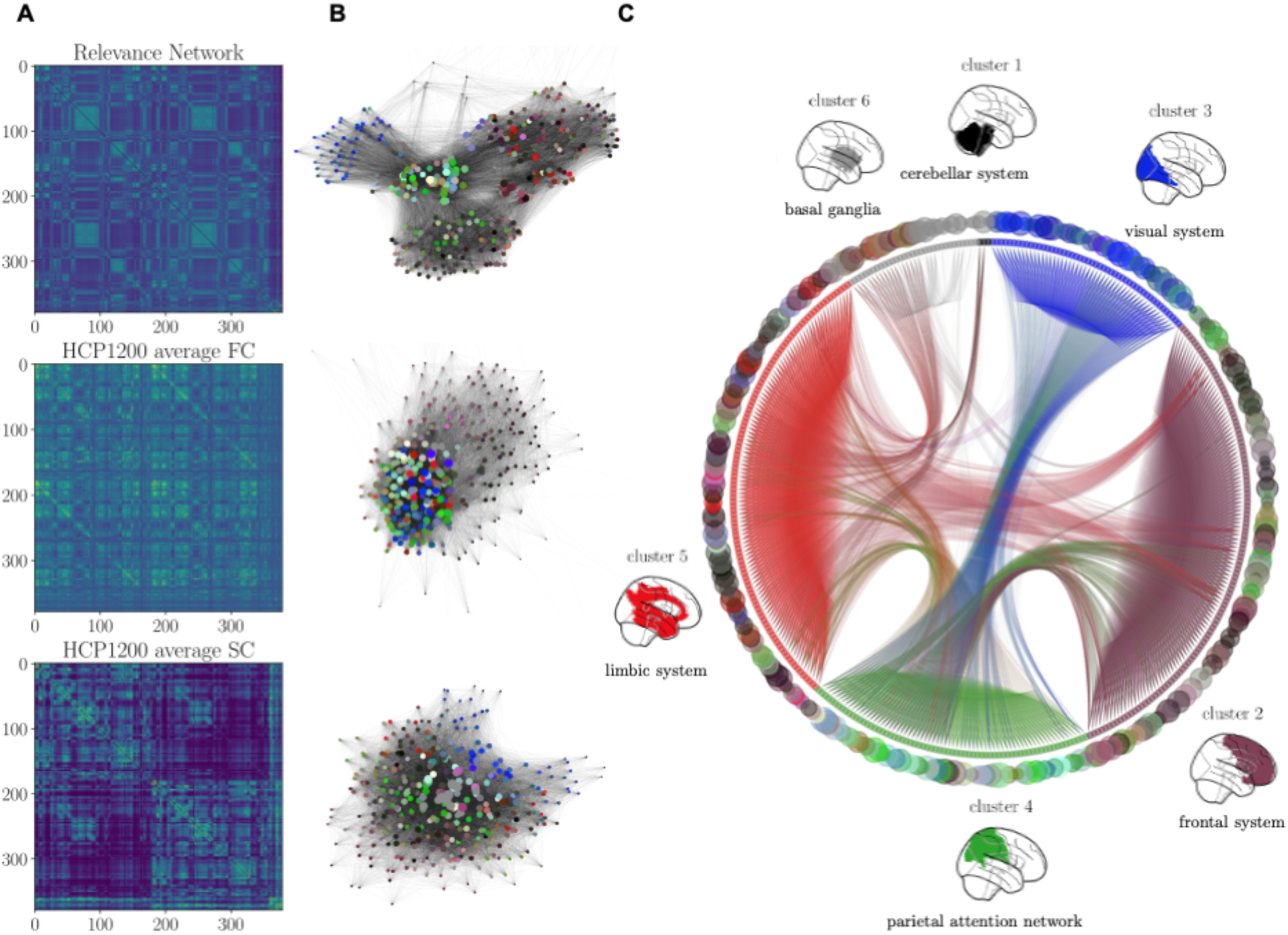
Relevance networks of co-occurring brain areas in TVBase maps resemble established functional and structural network patterns of the brain. A) shows a region-by-region matrix in Glasser parcellation, at the top depicting the correlation between relevance maps, in the middle the functional connectivity, and at the bottom the structural connectivity. It can be seen that the three modalities share particular patterns as the interhemispheric diagonals and distinct clusters of neighboring regions – which is observable because of the function- and anatomy-derived order of Glasser areas. B) visualizes the consecutive graph plots, including the color coding from Glasser 2016, describing five functional domains (task-positive in white, task-negative in black, auditory in red, visual in blue, and sensorimotor in green). C) Depicts six main clusters, separated by Louvain’s algorithm for community detection, showing similar topologies as known from neuroimaging data.

The correlations with both group-averaged SC (*r* = 0.31, *p* < 0.001) and FC (*r* = 0.27, *p* < 0.001) both indicate high similarity between the relevance network and empirical representations of brain structure and function. A multiple regression analysis shows that combined connectivity patterns explain more of the variance (*R^2^* = 0.132, p < 0.001) in the relevance network than each modality in separate. The SC has, hereby, a more prominent effect (*β* = 0.255, p < 0.001) than the FC (*β* = 0.186, *p* < 0.001).

A graph-based visualization of the networks (**Fig. 3B**) shows distinct characteristics of the three modalities, with the relevance network being most segregated into anatomically but also functionally distinct areas, here depicted by the color coding (Glasser, Coalson et al. 2016).

Hierarchical clustering of the relevance network highlights these results by resulting in six anatomically separate clusters (**Fig. 3C**). The specific pattern of inter-cluster associations indicates a stronger association between the visual system (cluster 3) and parietal attention network (cluster 4), as well as between the limbic system (cluster 5) and frontal areas (cluster 2). This highlights the potential of this novel modality to better understand interactions between different functional systems of the brain. As an example, this might apply to spatial orientation (visual and parietal) or emotion processing (frontal and limbic).

### 2.4 Symptom maps extend the interpretability of Amyloid PET and predict individual cognitive performance in Alzheimer’s Disease

To showcase the potential use of *TVBase* in translational settings, we investigated how it can help to interpret empirical neuroimaging data in Alzheimer’s Disease (AD). Amyloid PET is a powerful technique that is used in the diagnostic workup of cognitive decline as it maps Amyloid-beta, one of the hallmark proteins of AD. Besides the total and phosphorylated Tau protein, it is one of three diagnostic parameters in the current biomarker-based ATN classification of AD (Jack 2018). However, the interpretation of Amyloid PET still remains challenging and its broad use in clinical routine is controversially debated (Martinez, Vernooij et al. 2017, Jack, Bennett et al. 2018, Ruan and Sun 2023). Therefore, recent developments focus more on normalization techniques and spatial constraints, e.g., weighting particular brain regions more than others (Pemberton, Collij et al. 2022).

Hence, we explored the capability of *TVBase* to enrich Amyloid PET imaging by complementing it with symptom maps (**Fig 4A**). We used *TVBase* to map 37 AD symptoms as defined by the clinical criteria of the National Institute of Neurological and Communicative Disorders and Stroke and the Alzheimer’s Disease and Related Disorders Association (NINCDS-ADRDA, (McKhann, Drachman et al. 1984)). We correlated the spatial distribution of symptom maps with Amyloid PET data of 1127 subjects (552 female, mean age = 72 years) from the Alzheimer’s Disease Neuroimaging Initiative (ADNI, (Weiner, Veitch et al. 2017)). There are systematic differences between diagnostic groups, showing increased symptom-amyloid correlation with progressed disease stage from healthy controls to AD (**Fig. 4D**).

**Figure 4.**
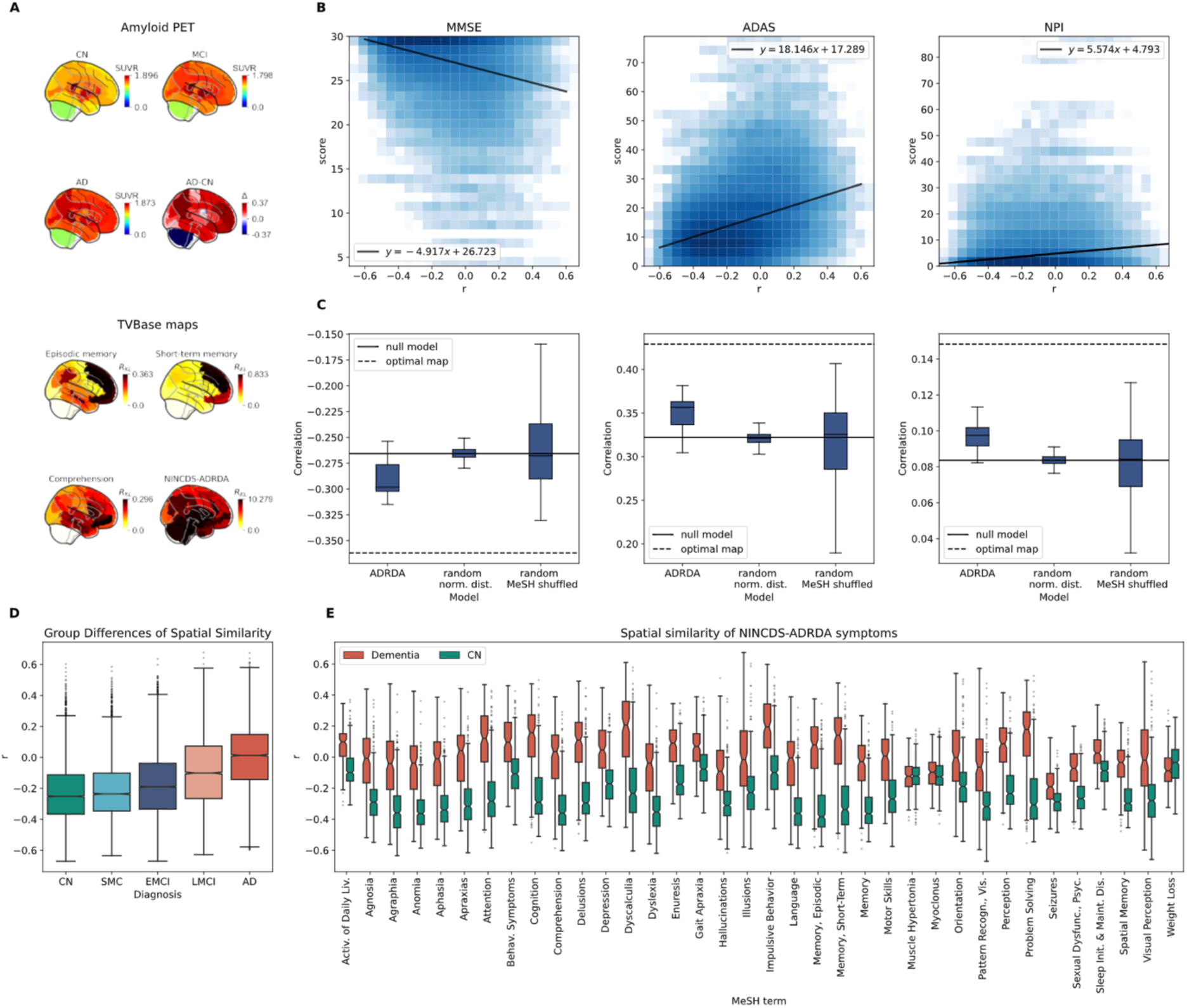
The similarity between Alzheimer’s Disease (AD) symptom maps and amyloid positron emission tomography (PET) with ^18^F-AV-45 supports the diagnostic classification and predicts cognitive scores. A) Average PET intensity for the diagnostic groups AD, mild cognitive impairment (MCI), and healthy controls (HC). Further the difference between AD and HC in the empirical data. Below are the three best-performing TVBase maps, “episodic memory", “short-term memory", and “comprehension", besides the average of all 37 TVBase symptom maps. B) There is a clear positive relationship between spatial similarity and symptom severity measured by neuropsychological scores with R^2^ ranging from 0.08 to 0.13. C) To assess the specificity of the correlation between whole-brain amyloid burden and each neuropsychological score, we first defined this as a null model for comparison. Further, an upper bound for the correlation – and therefore the predictive power – can be established by an optimal map. This optimal map is calculated by using a limited memory Broyden–Fletcher–Goldfarb–Shanno algorithm (Liu Dong and Jorge 1989) with 10-fold cross-validation. Finally, two types of random control maps are considered: one is sampled from a normal distribution based on the ADRDA maps, and the second is sampled from maps created out of all mapped MeSH terms. D) A mixed model ANOVA shows a significant main effect of spatial similarity for all diagnostic groups (η^2^ = 0.24) and all 37 symptom maps (η^2^ = 0.34) with strong effect sizes. E) Grouped boxplots showing the correlation between all 37 symptoms and amyloid PET of AD patients and healthy controls.

Further, this spatial similarity between symptom maps and Amyloid PET was used to predict the individual cognitive performance. Here, we used three standard neuropsychological assessments (**Fig. 4B**): the Mini Mental State Exam (MMSE), the Alzheimer’s Disease Assessment Scale – Cognitive Scale (ADAS-Cog), and the Neuropsychiatric Inventory (NPI). Weighting the amyloid burden with symptom maps significantly (*p* = 0.0079) improves the correlation with cognitive performance scores, compared to using empirical features alone, and to random *TVBase* maps (**Fig. 4C**).

Diving into the particular symptoms of AD, we show that almost all symptom maps differentiate between diagnostic groups by higher symptom-Amyloid correlation for AD patients than for controls (**Fig. 4E**). A pairwise t-test showed that similarity with brain maps of the terms “episodic memory”, “short-term memory” and “comprehension” differentiated best.

### 2.5 Decoding microscale pathways in Alzheimer’s Disease by augmenting TVBase with graph theory and Large Language Models

In the preceding sections, we have seen that TVBase maps contain information from empirical data and network characteristics and can be used in specific disease contexts. Nevertheless, the true strength of TVBase is that it is not limited to common neuroimaging data like fMRI or PET but also contains information from the microscale, e.g., genomic or post-mortem studies. Therefore, we aim to explore the microscale pathways involved in Alzheimer’s disease (AD) by combining TVBase with state-of-the-art technologies such as large language models (LLMs) and multi-layer graph analysis.

However, a key limitation of TVBase maps is their lack of directionality, which is, on the other hand, essential for understanding the flow at a molecular scale. To address this, we integrate LLMs to construct weighted TVBase maps, enhanced by an LLM-based assessment of the directionality of the association between the queried concept and the related brain regions (Fig 5B).

**Figure 5.**
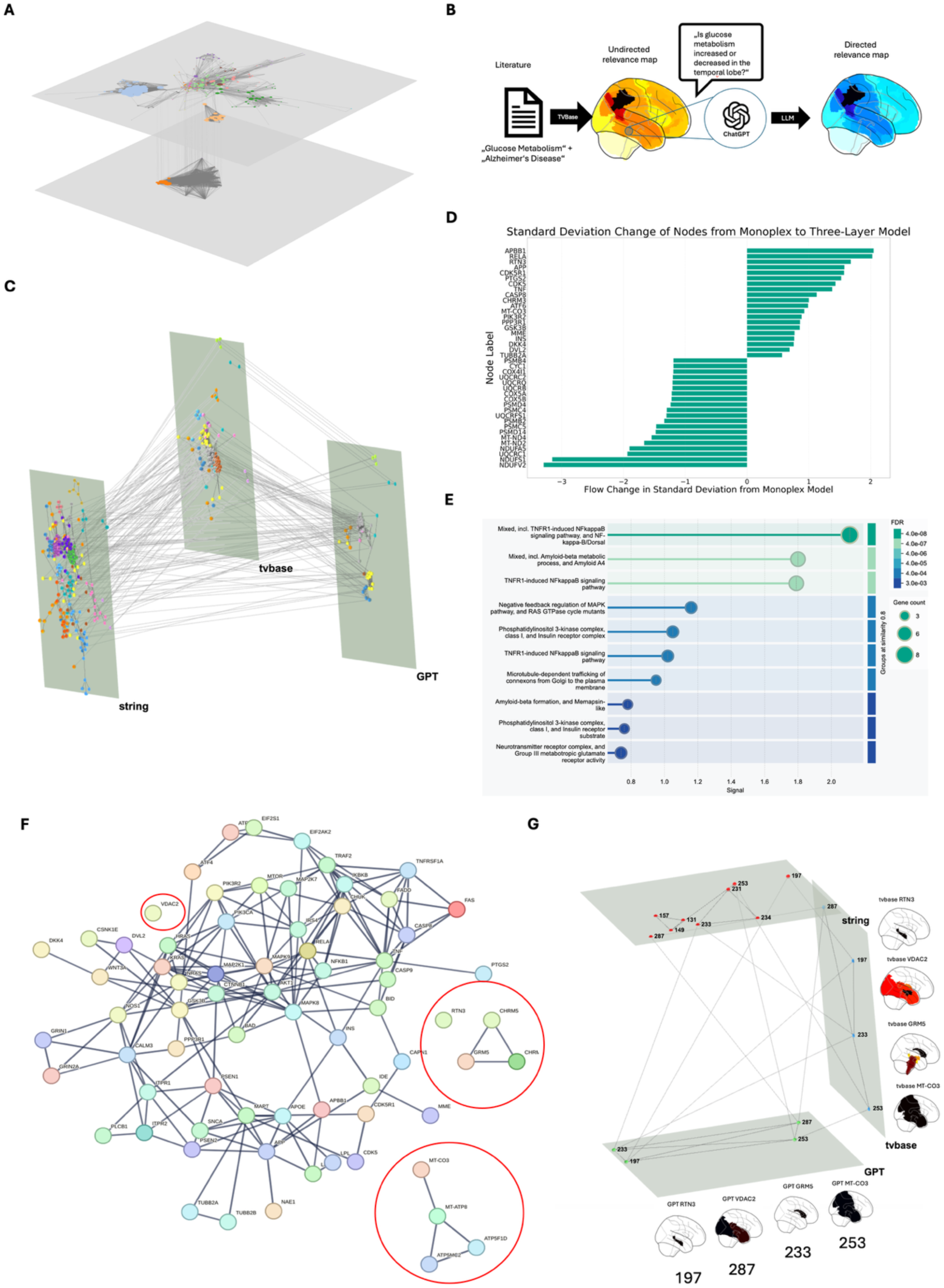
Exploring Alzheimer’s Disease (AD) pathways using multilayer-network analysis and LLM-augmented TVBase maps. A) To extend the semantic association within TVBase-maps by directionality, we employed the large language model (LLM) ChatGPT. The LLM rated each abstract used to assess whether the association between the Uberon term and the queried concept was positive, negative, or neutral, resulting in a weighted version of the original TVBase map. Node size corresponds to the information flow of that node B) Two-layer-network build of the TVBase maps for all 354 associated proteins in the KEGG pathway for AD: the protein-protein interactions from STRING build one layer, and the spatial similarity between TVBase maps based on their respective Pearson correlation builds the second. A flow analysis using the map equation (InfoMap) revealed stronger modularization but no change in the overall flow behavior. C) We constructed a third layer within our multilayer network depicting the directionalized similarity network of the LLM-enhanced maps. D) The information flow analysis showed a relevant shift within the network. E) Additional functional enrichment analysis showed a shift mostly towards inflammation-related proteins. F) The protein cluster with the highest flow in the three-layer network. Isolated nodes are highlighted, which can be considered knowledge gaps within the protein interaction network. G) The nine isolated nodes plotted in the three-layer network. Note that only four have non-empty TVBase maps, which are depicted beside the graph.

We start by utilizing the well-established Kyoto Encyclopedia of Genes and Genomes (KEGG) pathway for AD and represent it through the STRING protein-protein interaction (PPI) network, providing a possible ground truth for molecular relations. To facilitate the spatial context, we incorporate TVBase by constructing a network based on the spatial similarity between TVBase maps of proteins. These two layers are then considered to build a multi-layer network, in which each protein node is connected by separate edges based on the PPI network and the spatial similarity within TVBase maps and interlinked between the two layers. For the 351 proteins in the network, TVBase can add spatial information for 117 in the context of AD. The introduction of this additional layer led to different modular structures emphasized by a normalized-mutual information (NMI) score of 0.495 and adjusted rand index (ARI) score of 0.361. However, when analyzing the information flow, 93% of the model’s flow was still limited to the PPI layer (Fig 5A).

Thus, by creating a three-layer network (STRING, TVBase, and weighted TVBase), we tried to infer potential directional relationships between pathway components (Fig. 5C). This expanded model led to a more than three-fold increase in the information flow towards the spatial layers (30% vs 7%).

Remarkably, a flow analysis identified a shift in information flow (Fig 5D) towards nodes that are part of an inflammation module (Fig 5E). This protein cluster around TNF-alpha, NF-kappa-B signaling and Amyloid beta (Fig 5F) is therefore highlighted by the addition of spatial information and might be an area of particular interest in AD pathophysiology.

When analyzing this inflammation-related protein cluster (Fig. 5F), “islands” of disconnected nodes emerge. These “islands” are connected to the rest of the cluster via the two additional layers of the multi-layer graphs but not on the PPI layer of STRING, so they represent potential “knowledge gaps”. A closer examination of these knowledge gaps in three-dimensional space (Fig. 5G) alongside the corresponding weighted and unweighted TVBase maps suggests these four candidates (*RTN3, VDAC2, GRM5, MT-CO3*) for further investigating underlying mechanisms in AD.

## 3 DISCUSSION

In this article, we present the tool *TVBase*, an open-source software framework for aggregating and quantifying semantic knowledge about the brain preserved in the collective record of scientific literature. In other words, *TVBase* performs a semantic meta-analysis by calculating the association strength between an arbitrarily chosen biomedical concept and spatially distinct anatomical areas of the brain, which have been reported conjunctively in research articles. This is achieved by integrating literature mining and computational ontologies with state-of-the-art data standards and methods to analyze brain related data. The resulting brain maps rely on co-occurrences of queried terms and *Uberon* terms related to brain regions annotated in publication abstracts. These simultaneous citations can be used to reliably identify semantic relationships between these terms (Turney and Pantel 2010). To validate *TVBase*, we chose functional concepts from neuropsychology and cognitive neurology to systematically compare our results with already established and widely used tools for coordinate-based metanalysis and empirical neuroimaging data (R).

So far, *TVBase* bridges the fields of computational linguistics and neuroscience by projecting biological knowledge onto standard template brains, making it accessible and reusable for the neuroscience community.

The annotation with widely used terminologies and ontologies already decreases the ambiguity of language used in scientific literature. To further enrich the *TVBase* maps, it is possible to implement other knowledge representation systems about molecular pathways (Kanehisa, Furumichi et al. 2017, Martens, Ammar et al. 2021) or specific disease phenotypes (Iyappan, Gundel et al. 2016, Domingo-Fernandez, Kodamullil et al. 2017). Combining pathway databases with *TVBase* provides the unique opportunity to investigate where in the brain a particular cascade has been associated in the literature.

This knowledge can be transferred into computational models for whole-brain simulation. They can be constructed with the open-source neuroinformatic platform The Virtual Brain (TVB, www.thevirtualbrain.org), based on individual structural connectivity to simulate whole-brain activity (Ritter, Schirner et al. 2013, Sanz Leon, Knock et al. 2013, Sanz-Leon, Knock et al. 2015). TVB has been widely used in exploring dynamics in the healthy brain (Ritter, Schirner et al. 2013, Ritter, Born et al. 2015, Triebkorn, Zimmermann et al. 2020) and for modeling mental and neurological diseases (Zimmermann, Perry et al. 2018, Stefanovski, Triebkorn et al. 2019, Costa-Klein, Ettinger et al. 2020, Triebkorn, Stefanovski et al. 2022). By enriching brain network models with aggregated knowledge, the biological plausibility of these simulations can be further increased. Further, region-specific alterations of model parameters can be deduced from semantic association maps of brain disorders, which provides the potential to model spatial patterns of disease pathogenesis with biologically informed mechanistic network models.

Dynamical systems theories – as represented by TVB – inherit the potential to investigate the brain’s behavior *in silico*. Our software provides a robust computational algorithm to extract information from a human-incomprehensible amount of scientific research. However, ultimately, the final decisions must be made by the researcher. Therefore, many options for an in-depth review of the results are made available, including a per-paper review or specific filters for literature only referring to specific brain areas.

This eventually opens the door for personalized medicine applications, bridging individual data with common knowledge about the brain. So besides enriching simulations of common denominators of brain function, *TVBase* may also help identify biological mechanisms leading to brain structure changes or function in the individual, as it is even possible to project *TVBase* into the space of an individual brain to compare aggregated knowledge from millions of articles with data from one single subject.

Our newly developed tool, *TVBase*, poses the opportunity to interface with the semantic web and extract essential and multimodal data relevant to brain research. Further, our approach integrates a large quantity of semantic knowledge together with data from other databases into a common coordinate framework by transforming semantic concepts and their relations into brain coordinates. It is of striking necessity to represent and contextualize multimodal data across scales and build a comprehensive atlas of the human brain and its pathology (Rood, Stuart et al. 2019). Finally, a computational semantic framework like ours can act as an atlas of atlases linking several anatomical taxonomies – as the anatomical nomenclature has been developed “offline” for centuries before, resulting in a lack of concordance between anatomical descriptions (Bohland, Bokil et al. 2009).

*TVBase* transforms results from intensive literature mining into standard data frameworks for neuroscience, making means of information science accessible for applications like meta-analysis or brain simulation. Compared to other automated meta-analysis tools, the semantic association maps of *TVBase* are not limited to any specific method or fixed vocabulary. Hence it relies on the most extensive set of publications – almost all biomedical publications.

Conceptually, *TVBase* consequently relies on interoperability and integrability of knowledge. Integrating multimodal with other knowledge sources makes it even imaginable to generate cause-and-effect models of brain function and disorders that can be directly tested with TVB.

With concepts of open-source code and state-of-the-art data standards, *TVBase* provides a blueprint for future data technologies in neuroscience and enlightens the full potential promised by the computational processing of biological knowledge.

## 4 METHODS

This section describes the new methodology and the used data sources from a conceptual perspective. **Figure M1** gives an overview of the methodological pipeline of *TVBase*, from defining a biomedical concept and accessing knowledge sources to creating parcellated brain data. **Figure M1** gives an overview of the methodological pipeline. The complete pipeline description, including the code used, as well as the Supplementary, is currently available on request to the corresponding author.

**Figure M1.**
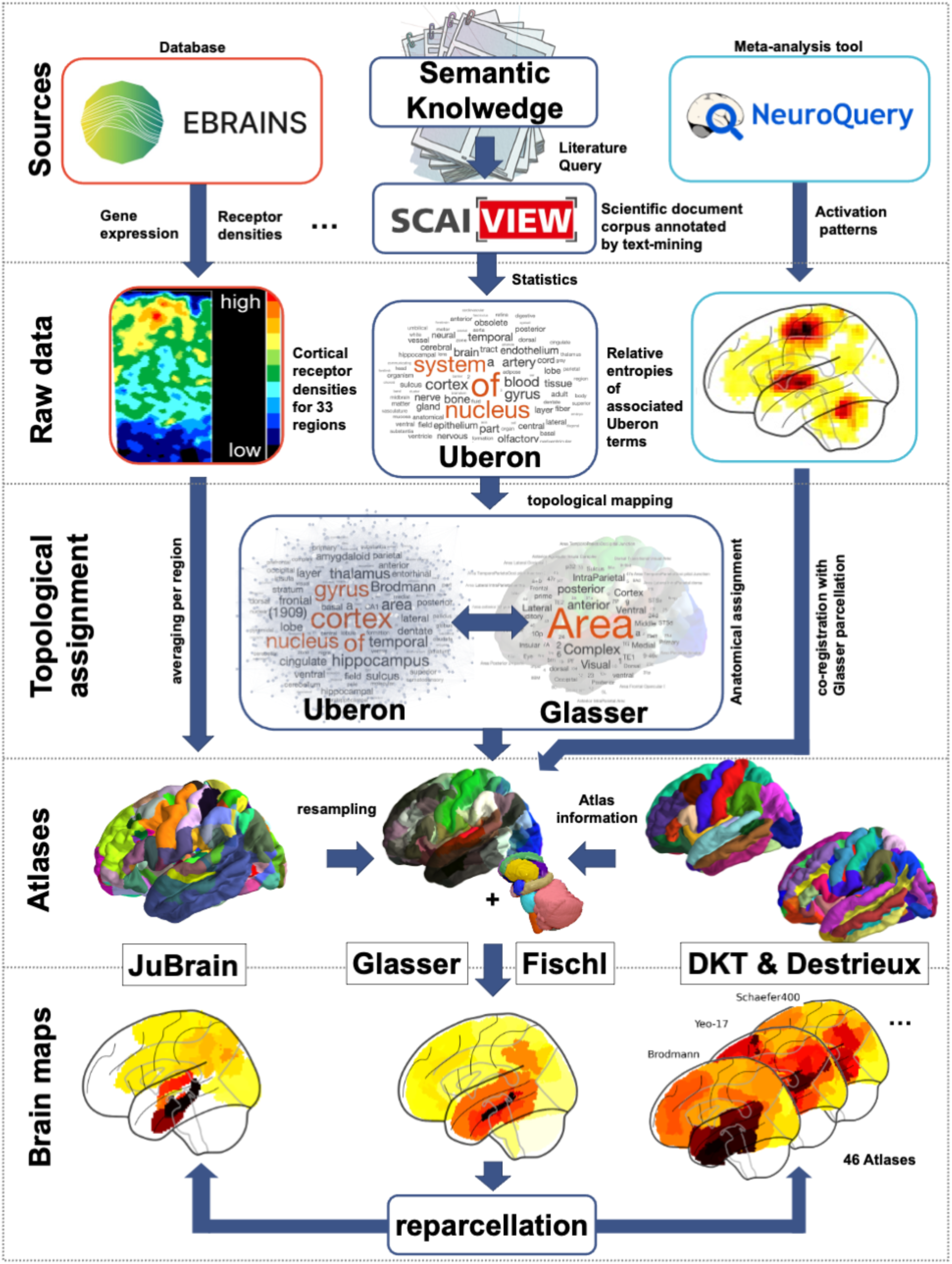
Workflow for the multi-source mapping methodology.

### 4.1 TVBase - Integrating Knowledge from Biomedical Literature into 3D Brain Space

*TVBase* is a software tool for mapping biomedical knowledge retrieved through literature mining onto a 3D template brain in standard reference coordinate space. While traditional meta-analyses usually condensate the underlying data of publications, a semantic meta-analysis refers to the semantic (meta-)information in the text document itself. Information from the scientific literature about brain areas involved or affected in specific biomedical processes – represented initially in natural language – is aggregated and transformed into a standard neuroimaging data format by the Neuroimaging Informatics Technology Initiative (NIfTI) (Larobina and Murino 2014). For an effortless integration with other brain research data, we chose the MNI-ICBM152 nonlinear brain stereotaxic registration model, version 2009c (further referred to as MNI-152) as template space (Fonov, Evans et al. 2009, Fonov, Evans et al. 2011).

To retrieve the annotated literature specific to a particular query, *TVBase* passes the user input of one or a more complex logical combination of search terms to the application programming interface (API) of the *SCAIView* software (https://api.academia.scaiview.com/). The query results are then again retrieved in the form of a specific literature corpus, listing all PubMed publications that entail all queried concepts in their abstracts. Additionally, all anatomical descriptions are extracted from this literature corpus defined in the detailed cross-species anatomy ontology *Uberon* (Mungall, Torniai et al. 2012). Consequently, by querying a particular concept or a complex combination of biological entities, *SCAIView* provides a comprehensive overview of all brain areas associated with that query in all PubMed publications based on co-occurrences at abstract level.

With this functionality, the association strength for each anatomical *Uberon* term specific to the literature corpus can be calculated by employing a measure of information theory: relative entropy. More precisely, a modified version of the Kullback-Leibler divergence (**Equation 1**) was calculated between the (discrete) Bernoulli distribution of the *Uberon* term’s frequency in the query literature corpus and its distribution in the total literature base (**Equation 2**). This divergence can be interpreted as measuring how frequently a brain region is mentioned in a particular publication set. This set consists of all publications matching the query compared to all publications in the literature. This highlights the region’s specificity to the given query and is therefore defined as relevance in the following. Consequently, *SCAIView* computes a relevance ranking of all anatomical *Uberon*-terms associated with a specific query.

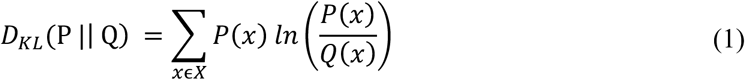

***Equation 1.** Original Kullback-Leibler Divergence compares two probability distributions, P and Q, for a continuous variable x.*

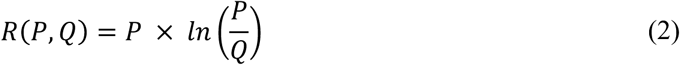

***Equation 2.** Modified Kullback-Leibler Divergence with P being the relative frequency of documents annotated with a term in the queried literature-corpus and Q being the relative frequency of documents with the term in the entire database.*

To relate these anatomical terms and their association strength to a common anatomical coordinate space, we carefully created a referential transformation matrix that maps the semantic anatomical classes of the *Uberon* anatomic ontology onto regions of a brain atlas. For a fine-grained representation of cortical areas, we mapped brain-related *Uberon* terms to the areas of the HCP multimodal parcellation (Glasser, Coalson et al. 2016), hereinafter referred to as Glasser parcellation. For an additional representation of subcortical areas, we used the modified Fischl subcortical segmentation from FreeSurfer (Fischl, Salat et al. 2002). For brevity, this combination of cortical and subcortical parcellations will be considered the *TVBase* atlas in the following.

To build the transformation matrix, we first extracted all 3853 *Uberon* terms annotated in *SCAIView*. Only those which unambiguously and hierarchically refer to the *Uberon*-term “Brain” were kept (e.g., a term like “kidney” cannot be spatially defined in the brain in a meaningful manner). From the initial anatomical terms of the *Uberon* ontology, 457 terms showed a distinct relation to brain anatomy. From those, 240 terms could be successfully mapped to at least one cortical area of the Glasser parcellation, and 183 were assigned to a subcortical region of a segmented template brain volume (Fischl, Salat et al. 2002). The remaining 34 terms were excluded, not referring to clearly defined cortical or subcortical areas. A detailed overview of inclusion criteria and exact mapping can be found in the anatomical revision section of the **Supplementary.** The transformation matrices are provided in **Supplementary Tables 2 and 3**.

Using this precise matching between *Uberon* terms and cortical Glasser areas and subcortical Fischl regions, *TVBase* projects the literature-mining results from *SCAIView* into a 3D coordinate system of the brain by adding the entropy-measure of each *Uberon*-term to its corresponding region of the template brain (**Equation 3**).

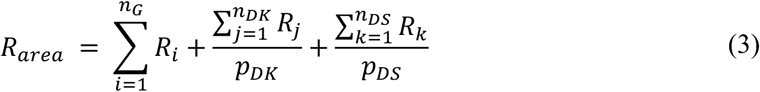

***Equation 3.** Cumulative relevance for each brain area, wherein R_area_ = area relevance score; R_i, j, k_ = relevance of Uberon terms, while i is referring to Glasser areas (G), j to areas from Desikan-Killiany (DK), and k to Destrieux areas (DS); n_G, DK, DS_ = the number of listed Uberon classes assigned to G or DK or DS. p= spatial overlap between DK/DS areas and corresponding Glasser areas.*

In the particular case of *SCAIView*, *P* and *Q* correspond to a single probability for each *Uberon* term in one query, with

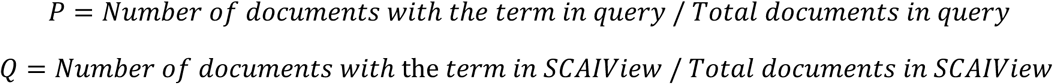

With this measure, relevance is high only if the *Uberon* term is mentioned disproportionately often in the given query, as opposed to the mere number of general occurrences. This is a special case of applicating the Kullback-Leibler divergence, as it formally does not fit the assumption that both distributions are defined over the same measurable space (as P is defined over the number of documents in one query, while Q is defined over the total number of documents). This leads to exceptional cases with (artificial) negative Kullback-Leibler divergence when the appearance of one term in a query is less frequent than expected a priori. This particular case is illustrated in **Figure M2**.

**Figure M2.**
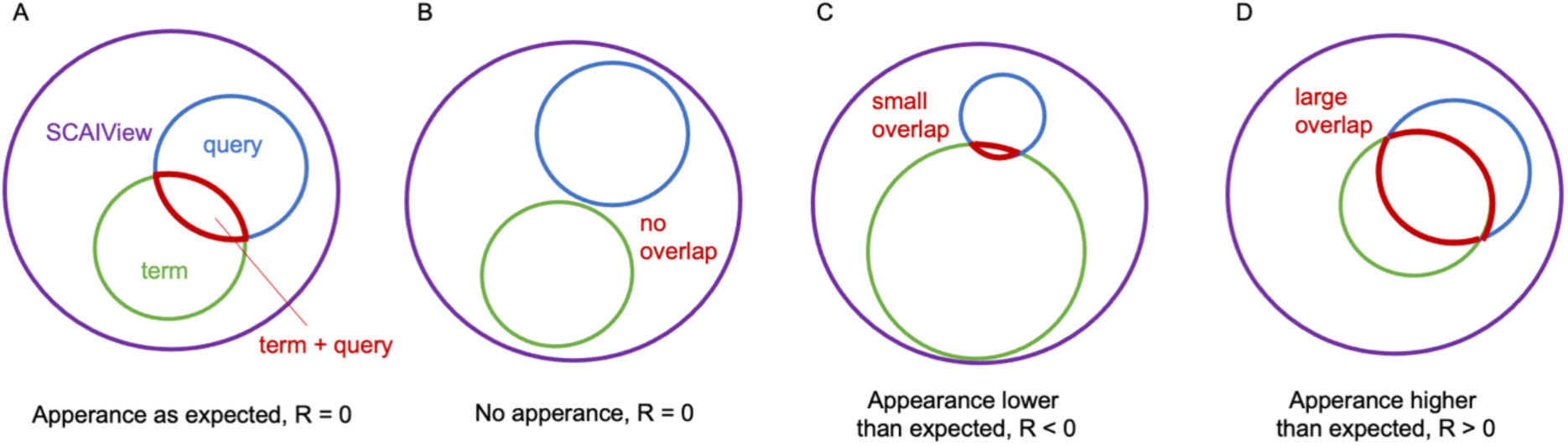
Special cases for relevance R, the modified Kullback-Leibler divergence (Equation 2). A) If the term is mentioned in the query as frequently as expected in the whole literature, the relevance is zero. B) The relevance is also zero for the special case of P = 0, as 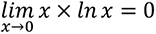 (L’Hôpital’s rule). C) The relevance is negative if the term is mentioned less frequently, as expected from the whole literature. Depending on the particular research question, a negative relevance can be ignored and therefore be set to zero. D) Only if the term is mentioned more often, as expected from the whole literature, do we have a real semantic association, and the relevance is higher than zero. Both a high occurrence of the term in the query and the rareness of the term in the rest of the literature can increase the relevance.

For P < Q, the relevance gets negative. As this introduces an artificial divergence inconsistent with the absence of the term (B), we set all negative relevance values to zero.

Meanwhile, D_KL_ is always positive, as can be proven by using Gibb’s inequality (Dragomir 2008). Therefore, the measure used in this work is an adapted Kullback-Leibler divergence between probabilities of different measurable spaces, which we will refer to as relevance in the following.

### 4.2 Interoperability between common brain atlases

Promoting a joint and flexible data representation system is necessary to integrate data from different brain atlases, like the *Multilevel Human Brain Atlas* from EBRAINS or the *Allen Human Brain Atlas*. To achieve atlas interoperability, we mapped the data arrays to widely used standard brain templates: The MNI152-ICBM2009c template and FreeSurfer’s *fsaverage* surface triangulation in MNI space. This allows us to “translate” the brain maps into other parcellations of the same template. The Glasser parcellation is a cortical surface parcellation, and regional data can be upsampled by assigning a unique region value to all vertices constituting that area in the surface triangulation. By overlaying another surface parcellation with the template brain *MeSH*, vertex data can be regrouped and averaged within the borders of one region from the new parcellation. This resampling between parcellations is analogically possible for volumetric parcellations. Therefore, we projected the group average surface of the Glasser parcellation to the volume of the MNI152-ICBM2009c template with segmented subcortical areas according to Fischl, Salat et al. (2002). We applied the same methodology as described above but based on voxel values.

This approach allows resampling the *TVBase* maps to all brain atlases registered to *fsaverage* or MNI space. While the original maps are parcellated according to Glasser (Glasser, Coalson et al. 2016), we used the parcellations of the cortical surface from *FreeSurfer* (Fischl 2012) to transform the maps into the atlases of Desikan-Killiany (Desikan, Segonne et al. 2006) and Destrieux (Destrieux, Fischl et al. 2010). Furthermore, we resampled our maps to another 40 different volumetric parcellations using MNI templates from (Lawrence, Bridgeford et al. 2021), available on https://github.com/neurodata/neuroparc, and to the detailed cytologic parcellation of the Julich-Brain (Amunts, Mohlberg et al. 2020), which we retrieved from EBRAINS.

**Table M1** provides an overview of all possible atlas transformations.

**Table M1.**
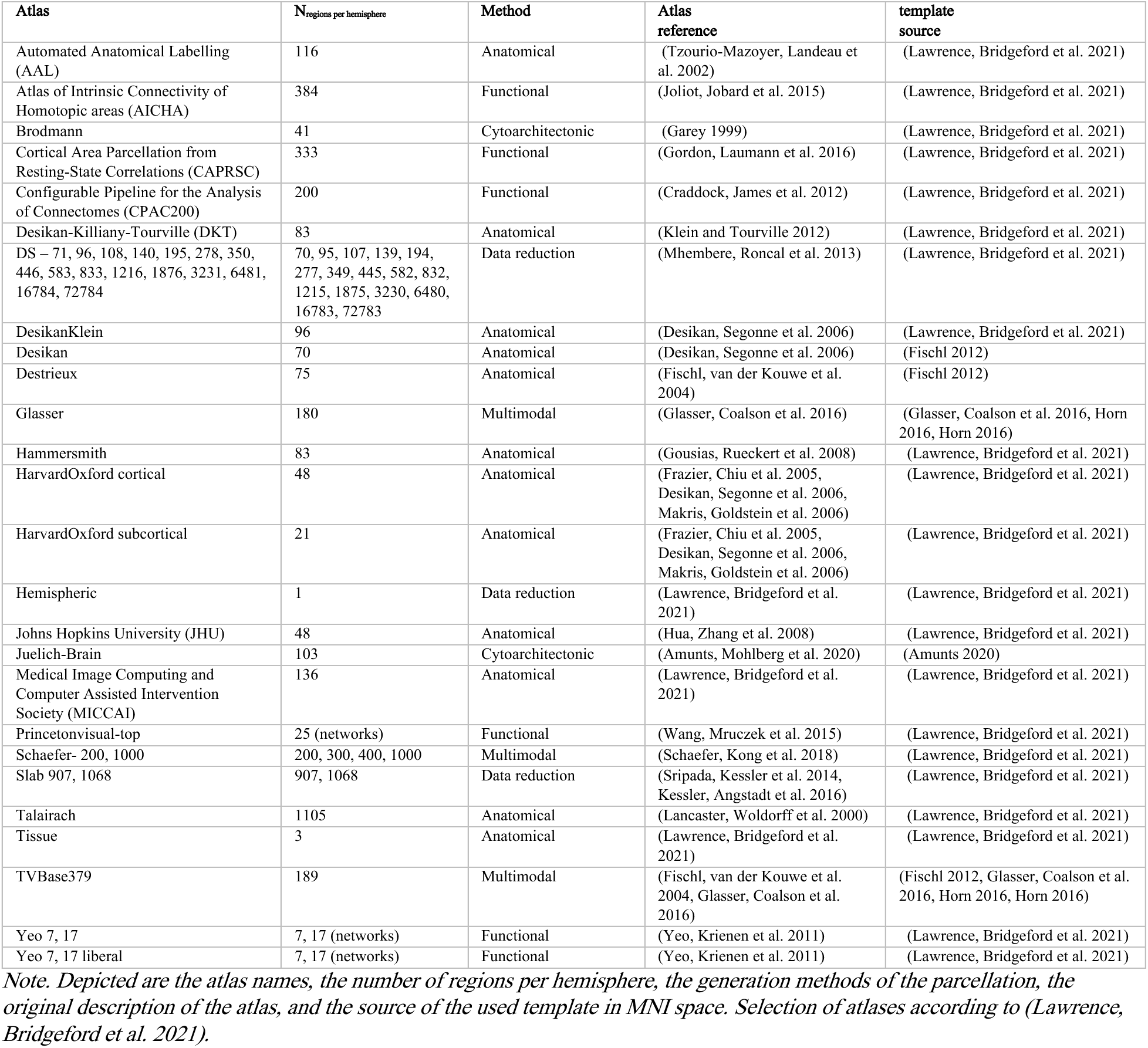
List of possible atlas transformations.

### 4.3 Validation

#### 4.3.1 Internal Validation

We mapped a broad range of terms from the Medical Subject Headings (*MeSH*) terminology to evaluate the performance of the proposed method. All papers of the MEDLINE database are indexed with the controlled vocabulary *MeSH* to facilitate detailed and comprehensive literature searches despite potentially coexisting or deviating descriptions of the same biomedical concept (Coletti and Bleich 2001). By using a *MeSH* term instead of a free-text query, a literature search is, on the one hand, more specific because it relies on a defined concept. On the other hand, the search is expanded with associated synonyms, making it independent of word choice. Hence, a detailed investigation of the “mappability” of different domains of the *MeSH* terminology contributes to the first assessment of possible fields of application. Therefore, three brain-related domains (i.e., trees) and sub-domains (i.e., branches) of the *MeSH* terminology will be systematically mapped using the proposed *TVBase* software and selectively reviewed. Mapping of the tree “Psychiatry and Psychology [F]”, as well as the branches “Central Nervous System Diseases [C10.228]” and “Brain Anatomy [A08.186.211]” will provide an extensive overview of the information aggregated with *TVBase*.

Moreover, mapping *MeSH* terms related to brain anatomy provides a recursive approach for validating the initial transformation matrix between *Uberon* and the Glasser parcellation. If the anatomical *MeSH* terms were correctly assigned to their corresponding areas in the 3D template brain, the accuracy of the anatomical transformation matrix could be confirmed.

By mapping the whole genome and all previously defined biological processes, we created a vast reference database for investigating data distributions of many maps.

#### 4.3.2 Systematic validation against automated coordinate-based meta-analysis tools

For further comparison and external validation, we implemented the recently published tool *NeuroQuery* (https://NeuroQuery.org) (Dockes, Poldrack et al. 2020), a tool for automated coordinate-based meta-analysis (CBMA) of brain imaging data. Like the functionalities in *TVBase*, *NeuroQuery* employs semantic literature-mining methods to associate a predefined vocabulary of relevant concepts with peak-activity coordinates reported in neuroimaging studies. Its database hosts brain coordinates of functional imaging data from 13459 publications, primarily from functional MRI studies. Building on that, *NeuroQuery* predicts a given query’s spatial activity pattern by assessing the searched term’s semantic similarities with stored concepts in the database. A linear regression then models the resulting brain map of spatial activity patterns associated with the queried concept (Dockes, Poldrack et al. 2020).

*NeuroQuery* offers an open-source Python package (https://github.com/NeuroQuery), which allows for the direct export of volumetric brain maps in *MNI-152* space for all concepts of the vocabulary. We resampled the data to fit the spatial resolution and orientation of the *TVBase* maps we created for each of the 7546 concepts. Then we systematically evaluated the overlap employing signal detection theory and calculating voxel-wise correlations in *TVBase* space.

The same comparison was run with maps from *Neurosynth (*https://neurosynth.org), another tool for creating automated CBMA results. Here, for each term in the *Neurosynth* vocabulary, we created subsets of studies and computed a coordinate-based activation likelihood estimation (ALE, (Eickhoff, Laird et al. 2009)) with a fixed sample size (*N* = 20). Consequently, the exact comparison measures were applied as we did for the validation with *NeuroQuery*.

### 4.4 Validation with empirical data from *neuromaps*

Further, we used multimodal brain maps, as published in *neuromaps* (Markello, Hansen et al. 2022), including various structural and functional imaging modalities. *Neuromaps* provides a data framework for accessing neuroimaging data from multiple sources in standard template spaces. Further, tools are available for interoperability between templates and projections from volumetric to surface data and vice-versa (Markello, Hansen et al. 2022). We used all 46 modalities from *neuromaps* that were queryable in *TVBase* (see **Table 1**).

To assess the similarity between the parcellated *TVBase* map and the empirical map, we calculated the voxel-wise Pearson correlation between both. We used two different permutation tests to assess this similarity’s significance further. First, we compared the similarity to random permutations of the same *TVBase* map. Second, we used a rotational shift of the data by iterating through all 379 indices of the *TVBase* parcels, similar to the voxel-wise approach described for *neuromaps* (Markello, Hansen et al. 2022). As a measure of specificity, we further randomly draw *TVBase* maps from the population of all *TVBase* maps for concepts from *MeSH,* for which we provide the percentile of randomly drawn *TVBase* maps that show a lower similarity to the empirical data than the semantically associated one.

### 4.5 Graph results methods

Minimally processed subject data from the HCP’s young adult S1200 release (Glasser, Sotiropoulos et al. 2013, Van Essen, Smith et al. 2013) was retrieved via DataLad, retrieved from: https://github.com/datalad-datasets/human-connectome-project-openaccess.

The Human Connectome Project was accessed on 27.04.2023 from https://registry.opendata.aws/hcp-openaccess.

Region-parcellation is based on HCP’s Glasser/MMP1 atlas (Glasser, Coalson et al. 2016), retrieved from: https://balsa.wustl.edu/file/87B9N.

The averaged FC matrix was created based on minimally preprocessed resting-state fMRI data from 1096 subjects (Glasser, Sotiropoulos et al. 2013). Individual functional connectivity matrices were calculated by a ROI-times-ROI correlation of the parcellated time-series from the denoised resting-state data in *fs_LR* space ("*rfMRI_REST <run>_<phase-encoding direction>_Atlas_MSMAll_hp2000_clean.dtseries.nii*”). FCs for each available run and phase-encoding direction were averaged separately for each individual and subsequently averaged across all 1097 subjects with available resting state fMRI data.

The structural connectivity was calculated based on anatomically constrained probabilistic tractography (Horbruegger, Loewe et al. 2019) using minimally preprocessed diffusion-weighted MRI data from 785 subjects (Glasser, Sotiropoulos et al. 2013). All subsequent analyses were performed using *MRtrix* (Tournier, Calamante et al. 2012) and FMRIB’s *FSL* (Jenkinson, Beckmann et al. 2012). First, a tissue-type segmented image was generated to differentiate between various tissue types, such as gray matter, white matter, cerebrospinal fluid, subcortical gray matter, and pathological regions. This segmented image was utilized to aid in the creation of a multi-shell, multi-tissue response function using *MRtrix’s* “dhollander” algorithm (Dhollander, Mito et al. 2019). Subsequently, a multi-shell, multi-tissue constrained spherical deconvolution algorithm (Tournier, Calamante et al. 2007) was employed to measure the fiber orientation distribution in each voxel. Based on these fiber orientations, full brain tractograms with approximately 25 million tracks per subject were created. Spherical-deconvolution-informed filtering of tractograms (SIFT) was applied to increase the biological plausibility of the fiber reconstructions (Smith, Tournier et al. 2013). The tracks were subsequently mapped onto a parcellated image to generate an ROI-times-ROI connectome. Since fiber-tracking was performed in the individual volumetric space, we registered the original Glasser parcellation to the individual surface space (via Freesurfer’s mri_surf2surf algorithm) and mapped it back to the volumetric T1w-image of the subject. Using Freesurfer’s automatic subcortical segmentation (Fischl, Salat et al. 2002) results, subcortical regions were added.

The TVBase relevance network (RN) was created based on 1186 parcellated brain maps from the *MeSH*-tree “Psychiatry and Psychology”, which we correlated across concepts to get a ROI-times-ROI correlation matrix of relevance. A multiple linear regression model with an interaction term (*RN* ∼ *SC x FC*) was used to assess the proportion of variance explained by either the structural connectivity or the functional connectivity. We thresholded the matrix at the 55th percentile for a graph-based visualization of the relevance network.

We performed hierarchical clustering (Bar-Joseph, Gifford et al. 2001) on the relevance matrix to group areas with similar data points. Condensed pairwise distances between the observations in the relevance matrix were used as input, which was calculated using the Euclidian distance between each pair of data points. The linkage method from the *SciPy* library was used to calculate the distance between the clusters with a farthest point algorithm. The linkage algorithm starts with a forest of *N* clusters, with *N* being the number of observations. In the following hierarchy, two clusters, *s* and *t*, are combined and removed from the forest. Instead, the combined cluster *v* is added. The algorithm terminates when only one cluster containing all observations is present. To get a reasonable number of clusters, the resulting dendrogram was cut at half the maximum distance between the clusters, providing six regional clusters. For all computations, we used the SciPy Python library (version 1.6.1).

### 4.6 Decoding data from the Alzheimer’s Disease Neuroimaging Initiative

Empirical data were obtained from the Alzheimer’s Disease Neuroimaging Initiative (ADNI) database (adni.loni.usc.edu). The ADNI was launched in 2003 as a public-private partnership led by Principal Investigator Michael W. Weiner. The primary goal of ADNI has been to test whether serial MRI, PET, other biological markers, and clinical and neuropsychological assessment can be combined to measure the progression of MCI and early AD. For up-to-date information, see www.adni-info.org. ADNI acknowledgments: http://adni.loni.usc.edu/wp-content/themes/freshnews-dev-v2/documents/policy/ADNI_Acknowledgement_List%205-29-18.pdf.

Post-processed and parcellated ^18^F-AV-45 (AV45) PET was retrieved from the ADNI database for 1127 subjects, including demographics and diagnoses. We selected standardized uptake values (SUVRs) for 91 subcortical and cortical grey matter areas from FreeSurfer’s Desikan-Killiany (DK) parcellation (Desikan, Ségonne et al. 2006). Further, data from three neuropsychological scores were accessed: The Mini-mental state examination (MMSE, (Folstein, Folstein et al. 1975)), the Alzheimer’s Disease Assessment Scale-Cognitive Subscale (ADAS-Cog, (Kueper, Speechley et al. 2018)), as well as the Neuropsychiatric Inventory (NPI, (Cummings, Mega et al. 1994)). Data at the base-line visit were available for five diagnostic groups, including subjects being cognitively “normal” (CN), having significant memory concern (SMC), with early and late mild cognitive impairment (EMCI, LMCI), as well as patients with Alzheimer’s Disease (AD).

To compare the individual Amyloid PET distributions with *TVBase* maps of AD symptoms, we mapped all 37 symptoms from the NINCDS-ADRDA classification system (McKhann, Drachman et al. 1984) and resampled them into DK, using *TVBase*’s reparcellation module. We also reparcellated 1102 *TVBase* maps of the *MeSH* trees “Psychiatry and Psychology” and “Neurological Diseases” into DK for comparison and specificity analyses. Using Spearman rank correlations, we assessed the spatial similarity between each subject’s spatial Amyloid deposition patterns and *TVBase* maps.

A one-way ANOVA was used to test group differences between diagnostic groups in the spatial similarity of all *TVBase* symptom maps with individual Amyloid PET. Further, we used a mixed-effects ANOVA with NINCDS-ADRDA symptom as within factors and the diagnostic group as between factors and the spatial similarity as a criterion to investigate which maps differentiate best the amyloid-PET patterns between diagnostic groups. Finally, general linear models were calculated to predict neuropsychological scores from the spatial similarity of the symptom maps with individual amyloid PET (Weiner, Veitch et al. 2017).

**Table M2.** Equivalent MeSH terms for the NINCDS-ADRDA catalog.

| NINCDS-ADRDA (McKhann 1984) | MeSH Term | MeSH ID |
| --- | --- | --- |
| memory | memory | D008568 |
| cognition | cognition | D003071 |
| language skill | language | D007802 |
| aphasia | Aphasia | D001037 |
| Perception | Perception | D010465 |
| visual perception | visual perception | D014796 |
| agnosia | agnosia | D000377 |
| Motor Skills | Motor Skills | D009048 |
| apraxia | Apraxias | D001072 |
| impaired activities of daily living | Activities of daily living | D000203 |
| altered patterns of behavior | Behavioral Symptoms | D001526 |
| depression | Depression | D003863 |
| insomnia | Sleep Initiation and Maintenance Disorders | D007319 |
| incontinence | Enuresis | D004775 |
| delusions | Delusions | D003702 |
| illusions | Illusions | D007088 |
| hallucinations | Hallucinations | D006212 |
| emotional, or physical outbursts | Impulsive Behavior | D007175 |
| sexual disorders | Sexual Dysfunction, Psychological | D020018 |
| weight loss | weight loss | D015431 |
| increased muscle tone | muscle hypertonia | D009122 |
| myoclonus | myoclonus | D009207 |
| gait disorder | Gait Apraxia | D020235 |
| seizures | seizures | D012640 |
| orientation to place and time | Orientation | D009949 |
| attention | attention | D001288 |
| problem-solving skills | problem solving | D011340 |
| social function | social interaction | D000083505 |
| calculation | Dyscalculia | D060705 |
| writing | Agraphia | D000381 |
| reading | Dyslexia | D004410 |
| ability to draw or copy | Pattern Recognition, Visual | D010364 |
| recent recall | Memory, Short-Term | D008570 |
| naming | Anomia | D000849 |
| repeating | see language |  |
| understanding | Comprehension | D032882 |
| spatial memory | spatial memory | D065852 |
| episodic memory | Memory, Episodic | D061212 |

### 4.7 Large Language Model weighted TVBase maps

OpenAI (www.openai.com) provides an API to access ChatGPTs capabilities, facilitating developers and researchers to integrate the model into their applications or research protocols. The primary method of interaction is through sending prompt-based queries to the API and receiving text-based responses.

*API Interaction Process:* for one of the here presented weighted TVBase maps we extracted all associated uberon-terms in the literature corpus and their corresponding abstracts. If there was no abstract present in the PubMed database, we referred to the title of the article instead. We then had a closer look to the association of the uberon term and context describing the content of the TVBase map (e.g. “Alpha rhythm”). To determine a potential directionality of the associations, we then made use of the ChatGPT API by sending a single shot **prompt** (text input to guide the model’s response). This prompt contained definitions of a potential positive association (e.g., “Alpha rhythm is *increased*”), as well as a corresponding negative association (e.g., “Alpha rhythm is *decreased*”). If the association was neither positive nor negative, or if there was no association to the context of the prompt at all, we defined this as a neutral association. The answer of the model was reduced to one word stating the most fitting definition: positive, negative or neutral

The specific prompt was as follows:

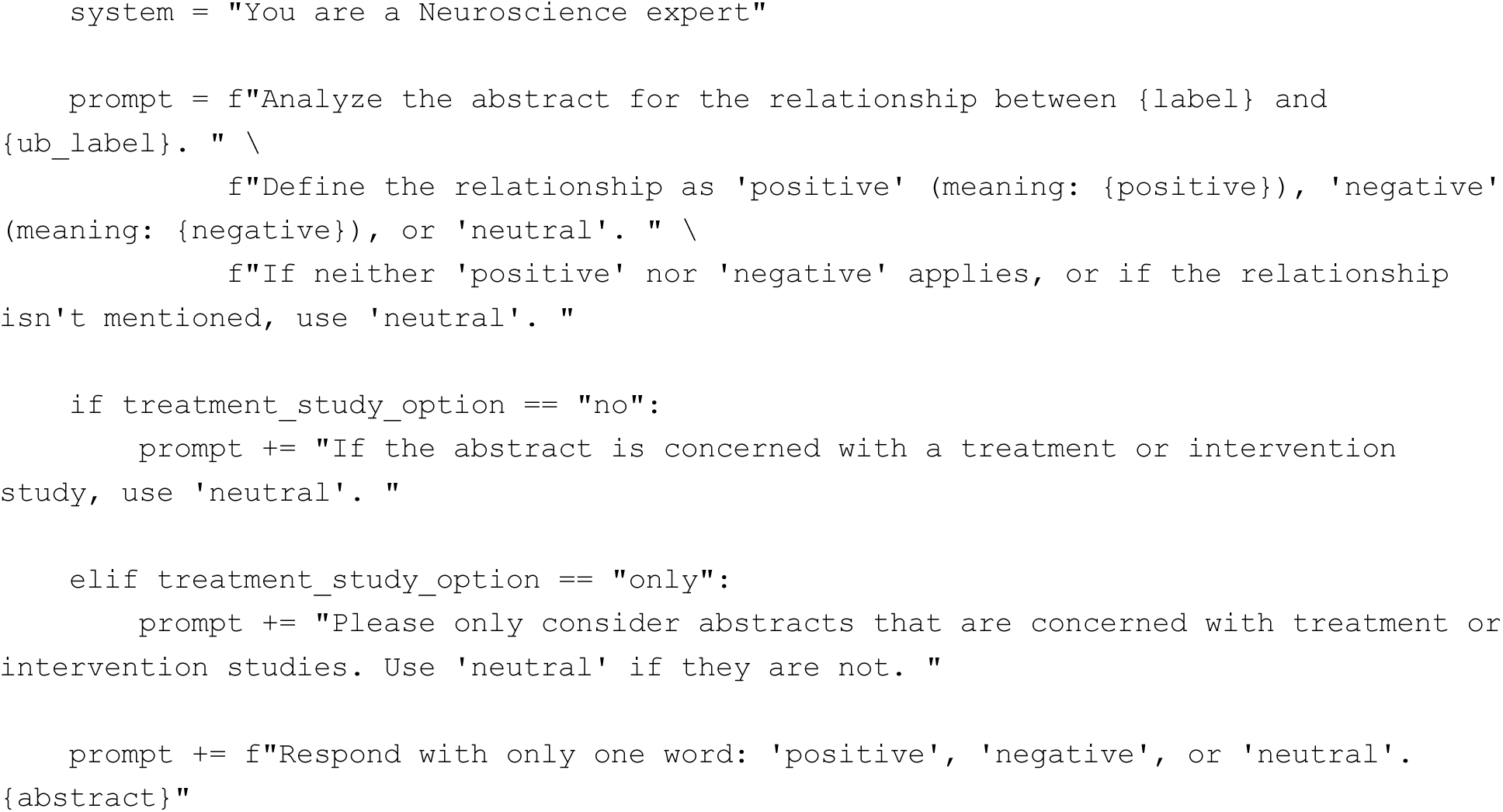

Where *label* corresponds to the label or context of the query (e.g. AD), *ub_label* corresponds to the brain region in question (e.g. temporal lobe), *positive* and *negative* refer to the definitions of directionality and *abs* corresponds to the abstract text.

The temperature parameter was set to 0.0 to have the maximum deterministic and coherent output.

The output was used to create a dictionary containing all relevant uberon terms, their corresponding Pubmed IDs (pmids), and their directionality according to our prompted definitions of directionality. The result was then used to weigh the relevance score of each uberon term in the given form:

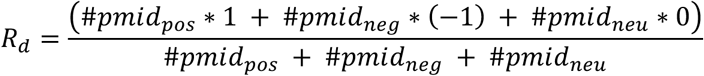

*R_d_* was then used to create new weighted TVBase maps.

We used the at the time of the creation of the results newest ChatGPT LLM version “gpt-4-1106-preview".

### 4.8 Multilayer network construction

To investigate the spatial organization of AD-related proteins, we constructed a multilayer network integrating protein-protein-interaction (PPI) data with spatial similarity metrics derived from TVBase and LLM-augmented TVBase maps. The first layer was built using the AD-specific PPI network from the Kyoto Encyclopedia of Genes and Genomes contained in STRING {Szklarczyk, 2023 #1096}, where nodes represented proteins and edges depicted experimentally validated interactions. The data was obtained via the STRING Rest-API (https://string-db.org/api/). We limited the output to include only interaction scores of the highest confidence level (evidence scores > 0.9). This led to a network containing 354 nodes with a density of 0.069, an average node degree of 24.3 and average local clustering coefficient of 0.654.

For each protein we then created a TVBase map by querying the HGNC term together with the MESH term for AD to extract the spatial information specific to AD. This led to 117 non-zero maps. These maps were then used to build a spatial similarity network by creating an adjacency matrix with the pandas library (version 2.1.1), in which elements represent the Pearson correlation between two maps. Therefore, each node represented the map of each protein, while the edges represented the spatial similarity. The spatial similarity network was then adapted to the same density as the PPI-network, leading to a threshold of 0.852 for the spatial similarity, an average node degree of 8.1 and a local average local clustering coefficient of 0.557.

For each protein TVBase map we created a LLM weighted map. These maps only contained the directional information of each semantic association, meaning that some maps were excluded, as they did not contain any clear directional information, or the negative and positive directionality cancelled each other out. This led to 88 weighted LLM maps. A spatial similarity graph was constructed in the same way as for the TVBase maps, leading to a threshold of 0.881 for the spatial similiarity, an average node degree of 6.1 and a local clustering coefficient of 0.524.

### 4.9 Network Analysis Using Infomap

We employed Infomap, a flow-based community detection algorithm, to compare network structures in monoplex and multilayer settings. The analysis was conducted in three configurations:

(1) a monoplex version consisting solely of the PPI network,
(2) a multilayer version incorporating both the PPI and spatial similarity layer from TVBase and
(3) a multilayer version incorporating the PPI and spatial similarity layers from TVBase and LLM-weighted TVBase maps.

Infomap optimizes the map equation to identify modules that maximize the flow retention within communities, thus revealing biologically meaningful clusters.

The edge weight is interpreted as flow between two nodes within InfoMap. This flow represents the probability of a random walker transitioning from one node to another. To account for the different biological intrerpretations of the weights within each layer and to make them more comparable in a multialyer configuration we used a couple of assumptions and data modifications:

As we only included PPI interactions with high confidence level > 0.9, the data showed a strong negative skew of -1.1175. After using a boxcox transformation this skewness reduced to -0.282. Spatial similarity on the TVBase layer works like repulsion and would mean that two proteins are less likely to act together. Within the LLM layer, the interpretation is more difficult, since strong negative correlation could mean that proteins act in the same areas, just in opposite directions. However, negative edge weights will be ignored in the map equation as they equate to a negative probability of a random walker passing along this edge. Hence, we only used positive spatial similarity on the TVBase layer and the absolute values for spatial similarity on the LLM layer.

To assess the influence of multilayer integration, we compared the steady-state flow of individual nodes between the mono- and multilayer cases. Node flow represents the probability of a random walker visiting each protein. Changes in node flow provided insight into how spatial organization refines functional associations beyond direct protein interactions.

The comparison of **node flow distributions** allowed us to determine whether certain proteins exhibited increased or decreased prominence when spatial information was integrated. Further, **modular overlap** and **reorganization** between the monolayer and multilayer clustering results were quantified using normalized mutual information (NMI) and adjusted Rand index (ARI) to evaluate the extent of similarity between detected communities.

All analyses were performed using the Infomap Python package, with parameters optimized for biological network applications.

The multilayer networks were visualized using the Python package pymnet and the Arena 3D web tool (Kokoli et al. 2023)

## Acknowledgements.

We acknowledge the assistance of ChatGPT, an AI language model developed by OpenAI, which was used solely for language refinement and proofreading purposes. All scientific content and interpretations remain the sole responsibility of the authors. The use of ChatGPT as a scientific tool as part of the results is independent of this and described in the Methods section.

We acknowledge support by EU Horizon Europe program Horizon EBRAINS2.0 (101147319), Virtual Brain Twin (101137289), EBRAINS-PREP 101079717, AISN 101057655, EBRAIN-Health 101058516, EIC grant PHRASE 101058240, by the Digital Europe Programme TEF-Health (101100700), Shaiped (101195135), CoordinaTEF (101168074), the German Research Foundation SFB 1436 (project ID 425899996); SFB 1315 (project ID 327654276); SFB 936 (project ID 178316478; SFB-TRR 295 (project ID 424778381); SPP Computational Connectomics RI 2073/6-1, RI 2073/10-2, RI 2073/9-1; DFG Clinical Research Group BECAUSE-Y 504745852, Berlin University Alliance OpenMake, the Virtual Research Environment at the Charité Berlin and EBRAINS Health Data Cloud and the Berlin Institute of Health and Foundation Charité.

Computation has been performed on the High-Performance Cluster for Research and Clinic of the Berlin Institute of Health, Berlin, Germany.

We thank Roopa Kalsank Pai for helpful discussions.

Data collection and sharing for the Alzheimer’s Disease Neuroimaging Initiative (ADNI) is funded by the National Institute on Aging (National Institutes of Health Grant U19AG024904). The grantee organization is the Northern California Institute for Research and Education. In the past, ADNI has also received funding from the National Institute of Biomedical Imaging and Bioengineering, the Canadian Institutes of Health Research, and private sector contributions through the Foundation for the National Institutes of Health (FNIH) including generous contributions from the following: AbbVie, Alzheimer’s Association; Alzheimer’s Drug Discovery Foundation; Araclon Biotech; BioClinica, Inc.; Biogen; Bristol- Myers Squibb Company; CereSpir, Inc.; Cogstate; Eisai Inc.; Elan Pharmaceuticals, Inc.; Eli Lilly and Company; EuroImmun; F. Hoffmann-La Roche Ltd and its affiliated company Genentech, Inc.; Fujirebio; GE Healthcare; IXICO Ltd.; Janssen Alzheimer Immunotherapy Research & Development, LLC.; Johnson & Johnson Pharmaceutical Research & Development LLC.; Lumosity; Lundbeck; Merck & Co., Inc.; Meso Scale Diagnostics, LLC.; NeuroRx Research; Neurotrack Technologies; Novartis Pharmaceuticals Corporation; Pfizer Inc.; Piramal Imaging; Servier; Takeda Pharmaceutical Company; and Transition Therapeutics.

